# GABAergic synapses between auditory efferent neurons and type II spiral ganglion afferent neurons in the mouse cochlea

**DOI:** 10.1101/2024.03.28.587185

**Authors:** Julia L. Bachman, Siân R. Kitcher, Lucas G. Vattino, Holly J. Beaulac, M. Grace Chaves, Israel Hernandez Rivera, Eleonora Katz, Carolina Wedemeyer, Catherine J.C. Weisz

## Abstract

Cochlear outer hair cells (OHCs) are electromotile and are implicated in mechanisms of amplification of responses to sound that enhance sound sensitivity and frequency tuning. They send information to the brain through glutamatergic synapses onto a small subpopulation of neurons of the ascending auditory nerve, the type II spiral ganglion neurons (SGNs). The OHC synapses onto type II SGNs are sparse and weak, suggesting that type II SGNs respond primarily to loud and possibly damaging levels of sound. OHCs also receive innervation from the brain through the medial olivocochlear (MOC) efferent neurons. MOC neurons are cholinergic yet exert an inhibitory effect on auditory function as they are coupled to alpha9/alpha10 nicotinic acetylcholine receptors (nAChRs) on OHCs, which leads to calcium influx that gates SK potassium channels. The net hyperpolarization exerted by this efferent synapse reduces OHC activity-evoked electromotility and is implicated in cochlear gain control, protection against acoustic trauma, and attention. MOC neurons also label for markers of gamma-aminobutyric acid (GABA) and GABA synthesis. GABA_B_ autoreceptor (GABA_B_R) activation by GABA released from MOC terminals has been demonstrated to reduce ACh release, confirming important negative feedback roles for GABA. However, the full complement of GABAergic activity in the cochlea is not currently understood, including the mechanisms that regulate GABA release from MOC axon terminals, whether GABA diffuses from MOC axon terminals to other postsynaptic cells, and the location and function of GABA_A_ receptors (GABA_A_Rs). Previous electron microscopy studies suggest that MOC neurons form contacts onto several other cell types in the cochlea, but whether these contacts form functional synapses, and what neurotransmitters are employed, are unknown. Here we use immunohistochemistry, optical neurotransmitter imaging and patch-clamp electrophysiology from hair cells, afferent dendrites, and efferent axons to demonstrate that in addition to presynaptic GABA_B_R autoreceptor activation, MOC efferent axon terminals release GABA onto type II SGN afferent dendrites with postsynaptic activity mediated by GABA_A_Rs. This synapse may have multiple roles including developmental regulation of cochlear innervation, fine tuning of OHC activity, or providing feedback to the brain about MOC and OHC activity.

**Significance Statement:** Cochlear OHCs receive efferent feedback from the brainstem to regulate auditory sensitivity and send afferent, feedforward information to the brain via type II SGNs. Histological evidence suggests an abundance of additional synaptic contacts in the OHC region, although neurotransmission at these synapses has not been determined. Here we demonstrate a synapse between efferent and afferent neurons that bypasses OHCs, and functions via GABA_A_R signaling. Although the function of this synapse is unknown, it is activity-dependent and persists in the mature cochlea, suggesting a role in auditory function.

## Introduction

Precise encoding of sound is mediated by powerful ribbon synapses between cochlear inner hair cells (IHCs) and type I spiral ganglion neuron (SGN) afferents. Contributions of cochlear outer hair cells (OHCs) are thought to occur via amplification of basilar membrane responses to frequency-specific sounds via prestin-mediated electromotility (Dallos and Corey, 1991; Oghalai, 2004; Ashmore, 2008; Dallos et al., 2008). OHCs also send information to the cochlear nucleus (CN) via glutamatergic synapses onto type II SGNs (Perkins and Morest, 1975; Ginzberg and Morest, 1983; Berglund and Ryugo, 1987; Simmons and Liberman, 1988; Benson and Brown, 2004; Weisz et al., 2009, 2021). Synapses between OHCs and type II SGNs are relatively sparse and weak compared to those between IHCs and type I SGNs (Weisz et al., 2012, 2014). These synaptic differences are thought to underly the higher sound thresholds and likely poor frequency tuning in type II SGN responses compared to type I SGN responses. Type II SGNs are also implicated in responses to cochlear damage (Liu et al., 2015; Nowak et al., 2021; Wood et al., 2021). The IHC and OHC regions also receive efferent innervation from the superior olivary complex (SOC) in the ventral brainstem. Within the SOC, lateral olivocochlear (LOC) neurons innervate peripheral processes of type I SGNs, while medial olivocochlear (MOC) neurons first form a transient developmental synapse onto IHCs (Glowatzki and Fuchs, 2000; Katz et al., 2004; Zorrilla de San Martin et al., 2010; Kearney et al., 2019), then form large cholinergic synapses onto OHCs (Warr and Guinan, 1979, reviewed in Guinan, 1996). The OHC response to acetylcholine (ACh) release from MOC neurons is mediated by alpha9/alpha10 nicotinic ACh receptors (nAChRs) that have a high calcium permeability. Calcium influx gates SK or BK potassium channels resulting in a larger, longer-lasting efflux of potassium (reviewed in Fuchs and Lauer, 2019). Therefore, the net effect of MOC activity on OHCs is hyperpolarization, which reduces OHC electromotility and thus reduces OHC-mediated contributions to the ‘cochlear amplifier’ (Galambos, 1956; Fex, 1962; Wiederhold and Kiang, 1970; Mountain, 1980; Siegel and Kim, 1982; Art et al., 1985). MOC inhibition of OHC activity is implicated in cochlear gain control, shifting the operating point of the cochlea between louder and softer sounds to allow cochlear responses to salient sounds, protection of the cochlea against noise-evoked trauma, and auditory attention (Oatman, 1976; Winslow and Sachs, 1987; Rajan, 1988; Kawase et al., 1993; Reiter and Liberman, 1995; Delano et al., 2007; Taranda et al., 2009; Maison et al., 2013; Terreros et al., 2016; Boero et al., 2018).

In addition to the canonical cochlear circuitry described above including between MOC efferent neurons and OHCs, and between OHCs and type II SGNs in the outer spiral bundle (OSB), additional synaptic connections have been described in histological and ultrastructural studies. These putative synapses include dendro-dendritic synapses between type II SGN dendrites, reciprocal synapses between OHCs and type II dendrites, synapses from the type II SGNs onto Hensen’s supporting cells, synaptic contacts between Deiters’ cells and afferent or efferent neurons, and direct synapses between MOC efferent neurons and type II SGN afferent neurons in the OSB (Eybalin et al., 1988; Burgess et al., 1997; Fechner et al., 1998, 2001; Thiers et al., 2002a, 2002b, 2008; Hua et al., 2021). More recent work also detected MOC efferent neuron synapses onto type I SGNs in the inner spiral bundle (ISB) (Hua et al., 2021). Despite supporting anatomical evidence, the developmental innervation patterns, neurotransmitter-mediated signaling, and ultimate function of these unique synaptic contacts have not been well-characterized.

In addition to ultrastructural analyses, histological studies in multiple species suggest that the inhibitory neurotransmitter gamma-aminobutyric acid (GABA) is synthesized in olivocochlear efferents. In particular, MOC efferent terminals have been shown to stain for the GABA synthetic enzyme glutamic acid decarboxylase (GAD), and for GABA itself (reviewed in Kitcher et al., 2021). Metabotropic GABA_B_ receptors (GABA_B_Rs) have been localized to MOC axon terminals as well as type I and II SGN somata and peripheral terminals (Maison et al., 2009; Wedemeyer et al., 2013). Several subunits of ionotropic GABA_A_ receptors (GABA_A_Rs) have also been localized to OHCs (Plinkert et al., 1989; Maison et al., 2006) and type I and II SGNs (Maison et al., 2006). At the single-cell level, GABA_B_R function has been demonstrated at MOC axon terminals, where GABA release activates pre-synaptic autoreceptors to decrease ACh release onto OHCs and developing IHCs (Wedemeyer et al., 2013). While there are reports that GABA affects OHC electromotility, others suggest no direct action of GABA on OHCs (reviewed in Kitcher et al., 2021). At the organismal level, GABA_A_R knockout mice exhibit hearing deficits and aberrant neuronal innervation (Maison et al., 2006). However, the localization of GABA_A_Rs on MOC somata in the brainstem confounds interpretation of these studies as systemic gene knockouts likely also impact central MOC function (Torres Cadenas et al., 2020). Thus, the role of GABA in the cochlea is incompletely understood.

The present study seeks to understand aspects of GABA signaling in the cochlea that remain unclear including whether GABA diffuses from efferent terminals in response to neuronal activity, and whether GABA_A_Rs function at synapses between MOC neurons and other cells. To establish GABA diffusion from MOC terminals to post-synaptic GABA_A_Rs in the cochlea, we combined patch-clamp electrophysiology with immunolabeling and optical detection of GABA. Using these methods, we were able localize both the site of GABA release in the OSB to MOC axon terminals, and the site of action of GABA_A_Rs to type II SGNs, which is consistent with previous ultrastructural studies detailing a synaptic contact between MOC efferent and type II SGN afferent neurons (Thiers et al., 2002b). We observe activity-driven release of GABA from MOC efferent axon terminals and confirm that the mechanism of GABA_B_ autoreceptor suppression of subsequent MOC neurotransmitter release is similar to that at the MOC-IHC synapse. Further, we demonstrate activity-dependent release of GABA from MOC axon terminals onto type II SGN afferent neurons that is established prior to hearing onset and persists in the mature cochlea in mice. While the function of this GABAergic MOC-type II SGN synapse is yet to be determined, interplay between pre-synaptic and post-synaptic effects of GABA signaling could have important roles in development of cochlear innervation, providing feedback to the brain about efferent activity, or fine-tuning of OHC activity.

## Methods

### Mice and ethical approval

Animal procedures followed National Institutes of Health guidelines, as approved by the National Institute of Neurological Disorders and Stroke/National Institute on Deafness and Other Communication Disorders Animal Care and Use Committee. Male and female mice were used in experiments. Mouse lines included wildtype (WT) C57BL/6J (Jax Cat No: 000664), ChAT-IRES-Cre (Jax Cat No: 028861), ChAT-IRES-Cre Δneo (Jax Cat No: 031661), Ngn-Cre^ERT2^ (Jax Cat No: 008529), Ai14 tdTomato reporter mice (Jax Cat No: 007914), and Bhlhb5-Cre (on Sv/129, C57BL/6J mixed background). Mice were housed on a 12/12 hr light/dark cycle, with continuous availability of food and water. Mice were anesthetized by carbon dioxide inhalation at a rate of 20% of anesthesia chamber volume per minute and then killed by decapitation.

### Immunohistochemistry

Mice aged postnatal day (P)5- 20 used for immunohistochemical experiments in cochlear whole mount preparations were euthanized as above. Temporal bones were immediately removed and fixed via perfusion of 1 ml cold 4% PFA (Electron Microscopy Science) in 1X PBS through the oval and round windows, with 20-30 min post-fix in cold 4% PFA. Cochleae were then washed with 1X PBS perfused through the round and oval window, then washed 3x 10-15 min in 1X PBS. Cochleae were either fully or partially dissected, followed by block of non-specific labeling (10% normal goat/donkey serum in 0.5% Triton X-100/1X PBS) for 1-2 hr, and washed for 15-20 min in 1X PBS. For experiments using an anti-GAD antibody (antibodies listed in **Table M1**), cochleae underwent an additional block in M.O.M. buffer (3-4 drops in 2.5 ml 1X PBS; Vector MKB-2213-1) at room temperature (RT) for 1 hr. After a 15–20-min wash, cochlear turns were incubated in primary antibody in blocking solution (Methods Table 1) at 4°C for 2-48 hr. Tissue was washed 3x 20 min in 0.25% TX-100 in 1X PBS, then incubated in secondary antibody in 5% normal goat/donkey serum and 0.25% TX-100 for 1-2 hr at RT. Tissue was washed for 15 min in 0.25% TX-100 in 1X PBS with DAPI (1:5000), washed 2x 10 min in 0.25% TX-100 in 1X PBS, then washed 15-20 min in 1X PBS.

**Table M1.**
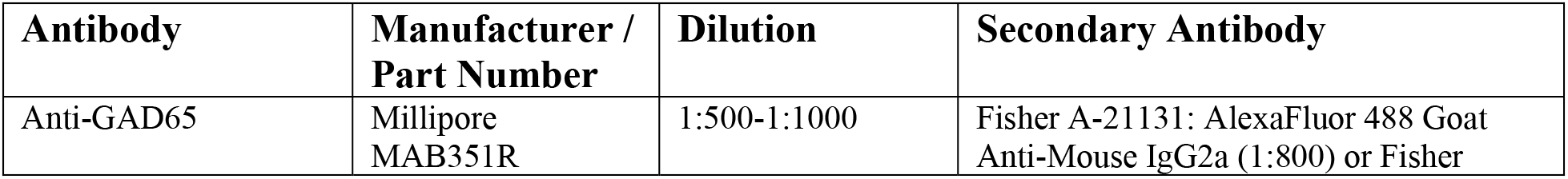
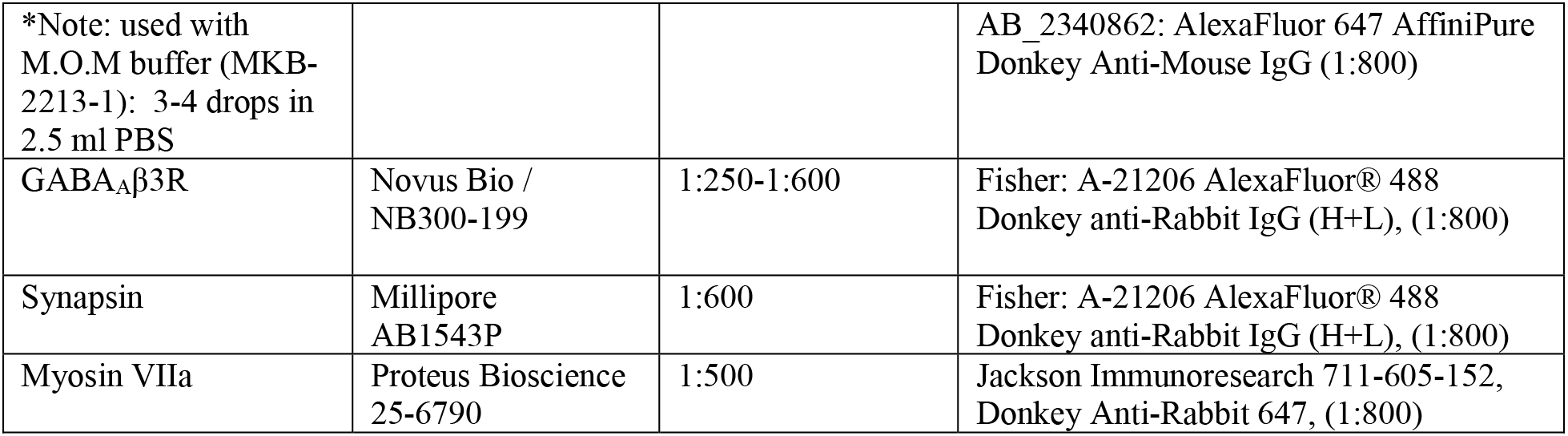
Antibodies used in immunohistochemistry experiments.

Antibody-labeled cochlear whole mounts were imaged on a Nikon A1R inverted microscope with 40X or 60X objectives, using Nikon Elements software v5.30.

### Patch-clamp recordings

For patch-clamp recordings from outer hair cells (OHCs), type II spiral ganglion neuron (SGN) afferent dendrites, and medial olivocochlear (MOC) efferent axons, the cochleae of P11-13 (OHC), P5-10 (type II SGN), and P13-17 (MOC) mice were dissected and placed under an insect pin affixed to a round glass coverslip with Sylgard (Dow Chemicals, Midland, MI, USA). Tissue was initially visualized using a Nikon Eclipse NI-E microscope with a QI-Click camera or FN-1 microscope with a pco.edge camera, with a 4X air objective for positioning of stimulation electrodes, drug application pipettes, or suction electrodes. Recordings were then performed using 40X or 60X water immersion objectives with DIC optics and an additional 1.25-2X magnification. For experiments with visualization of fluorescent cells, tissue was visualized under epifluorescent illumination using a Sola or Aura II lamp (Lumencor, Beaverton, OR, USA).

For OHC recordings, evoked IPSCs were obtained from the first row in the whole-cell voltage-clamp recording configuration with extracellular electrical stimulation of the MOC fibers. For type II SGN dendrite or MOC axon recordings, fibers under OHCs were exposed by removal of 4-6 OHCs using a glass pipette with a tip of 10-15 µm diameter. Patch-clamp recordings were performed using 1 mm diameter glass micropipettes (WPI) pulled (Sutter P1000) to resistances of 3-6 MΩ (OHC recordings) or 6-10 MΩ (type II SGN dendrite or MOC axon recordings). Dissection and recording ‘extracellular’ solution contained (in mM): 5.8 KCl; 155 (OHC recordings) or 150 (all else) NaCl; 0.9 MgCl_2_; 1.3 CaCl_2_; 0.7 NaH_2_PO_4_; 5.6 glucose; and 10 HEPES. The pH was 7.4, adjusted with 1N NaOH, and osmolarity = ∼315 mOsm. Extracellular solution at RT was perfused through the recording chamber at a rate of ∼1-2 ml/min. Internal solutions for type II SGN and MOC axon recordings contained (in mM): 50 KCl; 80 K-methanesulfonate; 5 MgCl_2_; 0.1 CaCl_2_; 5 EGTA; 5 HEPES; 5 Na_2_ATP; 0.3 Na_2_GTP; 5 Na_2_-phosphocreatine. The pH was adjusted to 7.2 with KOH. 0.01 mM AlexaFluor hydrazide 488 or 594 was added during some experiments to allow visualization of cell morphology to confirm neuron identity. Osmolarity was ∼290, liquid junction potential = -9 mV. Internal solutions for OHC recordings contained (in mM): 140 KCl, 3.5 MgCl_2_, 0.1 CaCl_2_, 5 EGTA, 5 HEPES, 2.5 Na_2_ATP. pH was adjusted to 7.2 with KOH, osmolarity = 283-290 mOsm. Drug application was performed using either a large bore gravity-fed glass pipette positioned close to the cochlear tissue (GABA or ACh application) or via addition to the re-circulating bath solution. Drugs were obtained from Sigma-Aldrich, Inc. (St. Louis, MO, USA), Alomone Labs (Jerusalem, Israel), and Thermo Fisher Scientific (Waltham, MA, USA).

Patch-clamp recordings were performed in voltage-clamp using a Multiclamp 700B amplifier with 1550B or 1322A Digidata controlled by Multiclamp Commander v11.2 and pClamp v11.2 (Molecular Devices, Silicon Valley, CA, USA) or a HEKA EPC 10 amplifier controlled by PatchMaster v2.91 (HEKA Instruments Inc.s, Holliston, MA, USA). Recordings were sampled at 50 kHz and lowpass filtered at 10 kHz. OHCs were held at -40 mV for recordings of MOC to OHC synaptic activity. Type II SGN and MOC axons were held at -80 mV for most recordings unless stated otherwise. To test the reversal potential of GABAergic currents, the type II SGN dendrite membrane potential was stepped to holding voltages from -80 to +20 mV in 20 mV increments for 10 seconds prior to GABA application. In all patch-clamp experiments, GABA application was performed once per 10 min.

### Electrical stimulation of MOC axons

Endogenous neurotransmitter release from MOC axons was evoked using electrical axon stimulation via glass micropipettes (Sutter Instrument Company, Novato, CA, USA) of ∼10-20 µm diameter containing extracellular solution described above, placed ∼20-30 µm modiolar to the inner spiral bundle (ISB). For OHC recordings, the stimulation electrode was positioned below the IHC aligned with the OHC under study. For optimal stimulation, cochlear supporting cells were gently removed by applying negative pressure through a glass pipette with the tip broken. The MOC axon stimulation was applied using an electrical stimulus isolation unit (Iso Stim, AMPI or SIU isolation unit A-M System). Stimulation timing and rate (for imaging experiments: 50 Hz, 1 second train duration) was controlled by the PClamp software during both patch-clamp and optical experiments.

### Posterior semi-circular canal (PSC) AAV injections

Posterior semicircular canal (PSC) injections to introduce AAV particles into the cochlea were performed as described in (Isgrig and Chien, 2018), using aseptic procedures. In brief, neonatal pups (P0-2) were hypothermia anaesthetized for ∼5 mins until they did not respond to stimulation, and then remained on an ice pack for the duration of the procedure. A postauricular incision was made using micro-scissors and the skin retracted. The PSC was identified under a surgical microscope and a glass micropipette pulled to a fine point was positioned using a micro-injector (WPI). For each mouse only one ear was injected with ∼1.2 µL AAV solution containing gene sequences encoding optical GABA indicator (iGABASnFR) variants (iGABASnFR.F102G, iGABASnFR2.0, or FLEX.iGABASnFR2.0). The incision was closed using a drop of surgical glue. About 4-5 pups per litter were injected. Pups were recovered to normal body temperature on a warming pad, while receiving manual stimulation to aid recovery. To increase the likelihood of the dam accepting the pups post-surgery, each pup was gently rubbed with bedding from the home cage and, if possible, urine collected from the dam using a cotton-tipped applicator, before being returned to their home cage. In addition, the dam had mineral oil applied to her nose to block detection of any surgical or human odors on the pups prior to being returned to the home cage.

### iGABASnFR imaging

For fluorescence imaging of iGABABSnFR-transduced cells in acutely dissected cochlear preparations, the euthanasia, dissections, extracellular solutions, drug application, and electrical stimulation of MOC axons were as above for patch-clamp recordings. iGABASnFR.F102G and FLEX.iGABASnFR.F102G were obtained as plasmids from Addgene.org and then packaged into AAV particles with the PHP.eB serotype and human synapsin (hSynap) promotor by Signagen. iGABASnFR2-containing plasmids were kindly gifted by the GENIE Project Team at Janelia Research Campus, Howard Hughes Medical Institute, and similarly packaged into AAV particles.

iGABASnFR imaging experiments were carried out using non-Cre-dependent virus injected into C57BL/6J mice or Cre-dependent (FLEX) virus injected into the Ngn1-CreERT2; tdTomato or Bhlhb5-Cre; tdTomato mice, which results in iGABASnFR expression specifically in neurons. The iGABASnFR or tdTomato fluorescence was localized to cells with the clear morphology of type II SGN dendrites. In experiments in which OHCs were also labeled, the focal plane of imaging was set to image only type II SGNs, or in the case of curved tissue that had focal planes containing both OHCs and type II SGN fibers, only the type II SGN-containing ROIs were analyzed (for analysis see *Image Processing*, below). Although unlikely, we cannot completely exclude the possibility that iGABASnFR was occasionally expressed in MOC neurons via AAV infection of CSF. However, analysis was restricted to regions with clear type II SGN morphology, and we did not detect expression in MOC axon terminals.

iGABASnFR imaging was performed on a Nikon A1R upright confocal microscope using resonant scanning in both red (568 nm, for tdTomato imaging in cochlear neurons from transgenic mice) and green (488 nm, for iGABASnFR variants) channels. In a subset of experiments with exogenous application of GABA or ACh, the ‘denoise’ function was utilized during imaging. In some experiments, a DIC-like image was simultaneously collected using the transmitted light detector, which converts the laser signal into a greyscale 3D image. Imaging settings included line averaging of 4-16 lines, bi-directional scanning, 512-1024 resolution, and frame rates of ∼7-15 frames per second.

### Imaging processing

Fluorescence intensity changes were measured in iGABASnFR-transduced neurons using ImageJ (NIH, Bethesda, MD, USA). A maximum intensity projection of the green (iGABASnFR) image stack was generated and then thresholded to set the regions to be used for region-of-interest (ROI) selection. Thresholds were set to 121, but manually adjusted in the case of tissue with brighter or dimmer fluorescence background (mean = 119 ± 20). The ‘analyze particles’ function was used to automatically draw ROIs around the iGABASnFR-expressing structures of interest, and each ROI was classified by eye as type I or type II SGNs from either fluorescence or transmitted detector images. It was not possible to identify individual neurons from either tdTomato or iGABASnFR images, so ROIs likely contain multiple neuron segments. These ROIs were then used to measure fluorescence intensities in the original image stack for each frame. Fluorescence intensity values per ROI and per frame were imported into Origin v2021 (Origin Lab, Northhampton, MA, USA). The baseline mean and standard deviation of fluorescence was measured for one second prior to stimulus onset (electrical MOC axon stimulation) or ten seconds prior to drug application (exogenous GABA or ACh application). The ΔF/F of iGABASnFR responses was determined for the 30 seconds of exogenous ACh or GABA application compared to the 10 seconds prior to drug application. For experiments utilizing electrical stimulation of neurotransmitter release from MOC axons, first we determined whether each ROI had a positive ‘response’ to the axon stimulation, defined here as a maximum fluorescence greater than twice the mean of the baseline plus two standard deviations of the baseline (mean +2SDs). To prevent a noisy fluorescence signal from giving an artificially high ‘maximum’ intensity, we used a rolling average (5 frames before, 5 frames after) to smooth the trace, and determined the fluorescence maximum from this rolling average trace. We then calculated the mean and standard deviation from 1 sec prior to stimulation in the non-averaged trace. The mean +2SDs was subtracted from the ‘maximum’ to detect positive values that were categorized as a ‘response’. To determine the ΔF/F of the fluorescence following MOC axon stimulation, the mean of the baseline fluorescence was subtracted from the maximum fluorescence, then divided by the mean baseline fluorescence.

### Statistics and Software

Statistics were performed in Origin v2021-2023 or GraphPad Prism 6 (RRID:SCR_002798). Data were tested for normality using a Shapiro-Wilk test. Statistical tests included a one-way ANOVA for parametric tests. For non-parametric statistical analyses, with two groups, a Wilcoxon rank sum was applied, and for non-parametric comparisons of more than two groups, a Friedman test followed by a Dunn multiple-comparison test was used. Figures were prepared in Origin, ImageJ, and Adobe Illustrator (Adobe, San Jose, CA, USA).

## Results

### Localization of GABAergic pre-synaptic structures in the cochlea

Previous work has shown immunolabeling for the GABA synthetic-enzyme GAD in cochlear structures with the typical morphology of LOC and MOC efferent neurons (Fex et al., 1986; Maison et al., 2003). We first confirmed GAD immunolabeling in olivocochlear terminals of cochlear preparations from ChAT-IRES-Cre; tdT mice, which have been previously characterized for genetic identification of MOC neurons (Torres Cadenas et al., 2020; Romero and Trussell, 2021). GAD-immuno-positive structures were clearly visible by postnatal day 7 (P7, **Figure 1A-E**) and attained the typical pattern of olivocochlear innervation of the OSB immediately prior to the onset of hearing, between P9 (**Figure 1F-J**) and P11 (**Figure 1K-O**). No apparent change in labeling was observed immediately after hearing onset from P13 (**Figure 1P-T**) through at least P20 (**Figure 1U-X**). In addition, GAD-positive MOC axon terminals from ChAT-IRES-Cre; tdT mice also colocalized with the presynaptic marker Synapsin, suggesting activity-dependent neurotransmitter release (**Figure 1U-X**). Taken together, early labeling of efferent terminals with GAD indicates that MOC neurons synthesize GABA during postnatal cochlear development, while labeling at later ages suggests GABAergic signaling is a stable feature of the mature, hearing animal.

**Figure 1.**
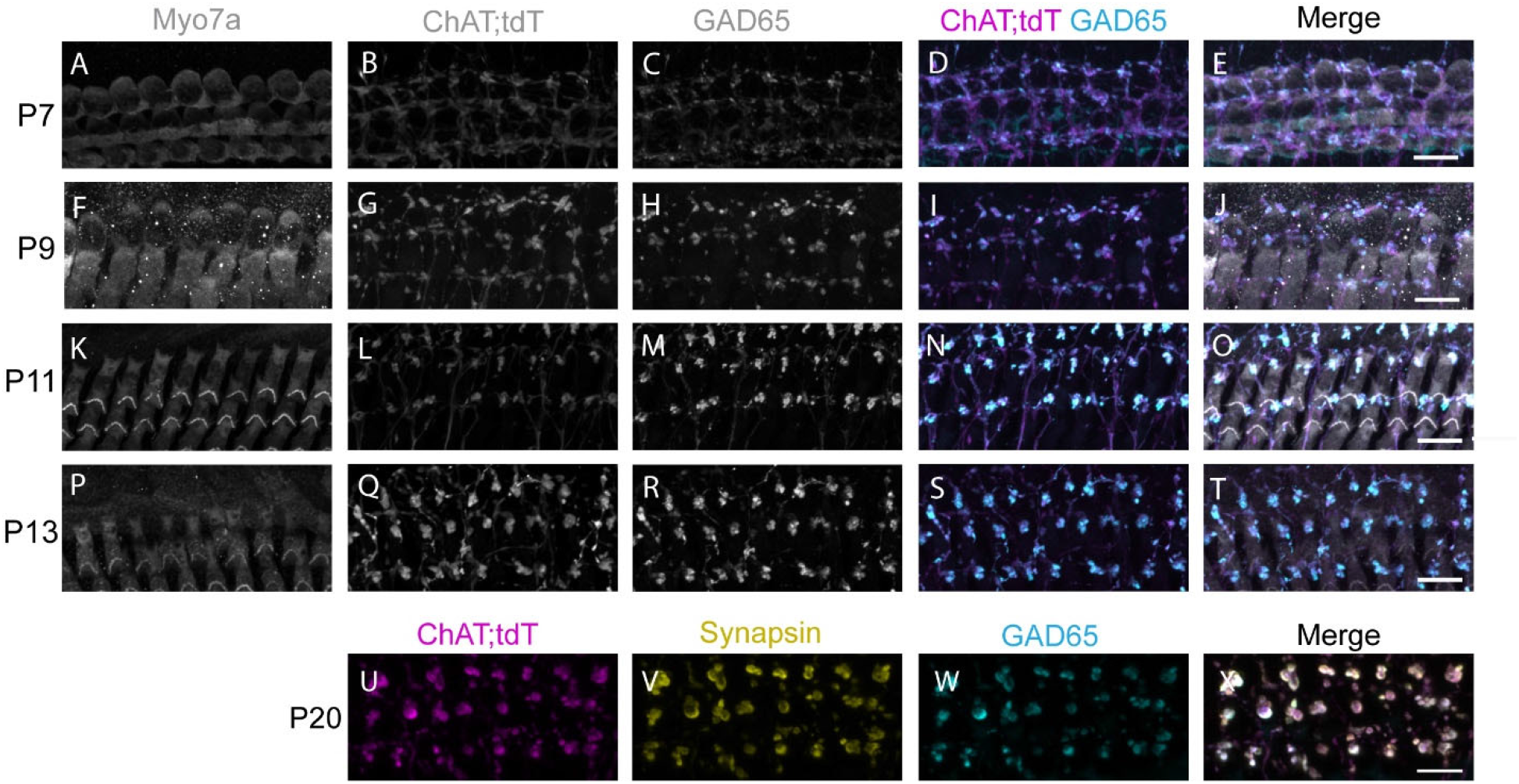
Genetically identified MOC axon terminals are GABAergic. A-T. Immunolabeling in ChAT-IRES-Cre; tdT mouse medial turns at the ages indicated. A,F,K,P: Immunolabeling for Myosin VIIa (greyscale) labels OHCs. B,G,L,Q: ChAT-IRES-Cre; tdT (ChAT; tdT, greyscale) signal labels MOC efferent axons. C,H,M,R: GAD65 (greyscale) labels puncta under OHCs. D,I,N,S: Merge of ChAT-IRES-Cre; tdT (magenta) and GAD (cyan) shows colocalization of the two channels. E,J,O,T: Merge of all channels. U-X. Immunolabeling in ChAT-IRES-Cre; tdT mouse apical turn at P20. U. ChAT-IRES-Cre; tdT (ChAT; tdT, magenta) signal co-localizes with V. Synapsin (yellow) and W. GAD (cyan). X. Merge of U-W. Scale bar in all panels is 10 µm.

### GABA reduces MOC neurotransmitter release via presynaptic GABA_B_Rs

To confirm the release of GABA from MOC axons in the OSB and explore the release mechanism, we first performed recordings from cochlear OHCs in P11-13 mice while electrically stimulating MOC axons to evoke neurotransmitter release (**Figure 2A**). Consistent with previous results in mouse OHCs from cochlear apical turns, MOC axon stimulation evoked post-synaptic currents that are mixed nAChR and SK potassium currents (Wersinger et al., 2010; Ballestero et al., 2011; Wedemeyer et al., 2013; Vattino et al., 2020) (**Figure 2B**). In previous work at this synapse, it was shown that GABA negatively modulates the amount of transmitter released from MOC neurons by activating presynaptic GABA_B_Rs (Wedemeyer et al., 2013). Therefore, we first evaluated the effect of the GABA_B_R antagonist CGP 36216 (100µM) (**Figure 2B**). Our results are consistent with those reported by Wedemeyer et al., 2013, and clearly show that there is a significant increase in the quantal content of transmitter release in the presence of this antagonist (control: 0.15 ± 0.03, CGP: 0.41 ± 0.08 Wilcoxon rank sum test, p = 0.0313, n = 6 cells). At P11-13, transmitter release at the MOC-OHC synapse is supported by Ca^2+^ entry through P/Q- and R-type voltage gated calcium channels (VGCCs) (Vattino et al., 2020). We therefore evaluated whether the activation of GABA_B_Rs was interfering with Ca^2+^ entry through either or both channels. As illustrated in **Figure 2C**, when blocking P/Q-type VGCCs with ω- agatoxin IVA (200 nM), there was a significant reduction in the quantal content of release. Under this condition, application of the GABA_B_R antagonist CGP caused no additional effect on neurotransmitter release (control: 0.20 ± 0.04, ω-agatoxin: 0.06 ± 0.02; ω-agatoxin + CGP: 0.05 ± 0.01. Friedman test (with post hoc Dunn’s test), control vs. ω-agatoxin, p = 0.0177, control vs. ω-agatoxin + CGP, p = 0.0177, ω-agatoxin vs. ω-agatoxin + CGP, p = 0.92 (NS); n = 5 cells.). However, when blocking R-type VGCCs with Ni^2+^ (100 µM), a strong blocker of this type of VGCC (Myoga and Regehr, 2011; Vattino et al., 2020), further application of CGP 36216 still caused a significant increase in the quantal content of release, indicating that the effects of GABA_B_Rs are not mediated by inhibition of R-type VGCC (control: 0.22 ± 0.03, Ni^2+^: 0.10 ± 0.0, Ni^2+^+CGP p = 0.24 ± 0.03. Friedman test (with post hoc Dunn’s test), control vs. Ni^2+^ p = 0.0269, control vs. Ni^2+^ + CGP p = 0.75 (NS), Ni^2+^ vs. Ni^2+^ + CGP p = 0.0114, n = 5) (**Figure 2D**). These results are consistent with those shown at the transient MOC-IHC synapse from P9-11 mice (Wedemeyer et al., 2013). In conclusion, the present results suggest that GABA released by MOC OSB axons is regulating both ACh and potentially its own release by acting on presynaptic GABA_B_Rs that in turn reduce the amount of Ca^2+^ flowing through P/Q-type VGCCs.

**Figure 2.**
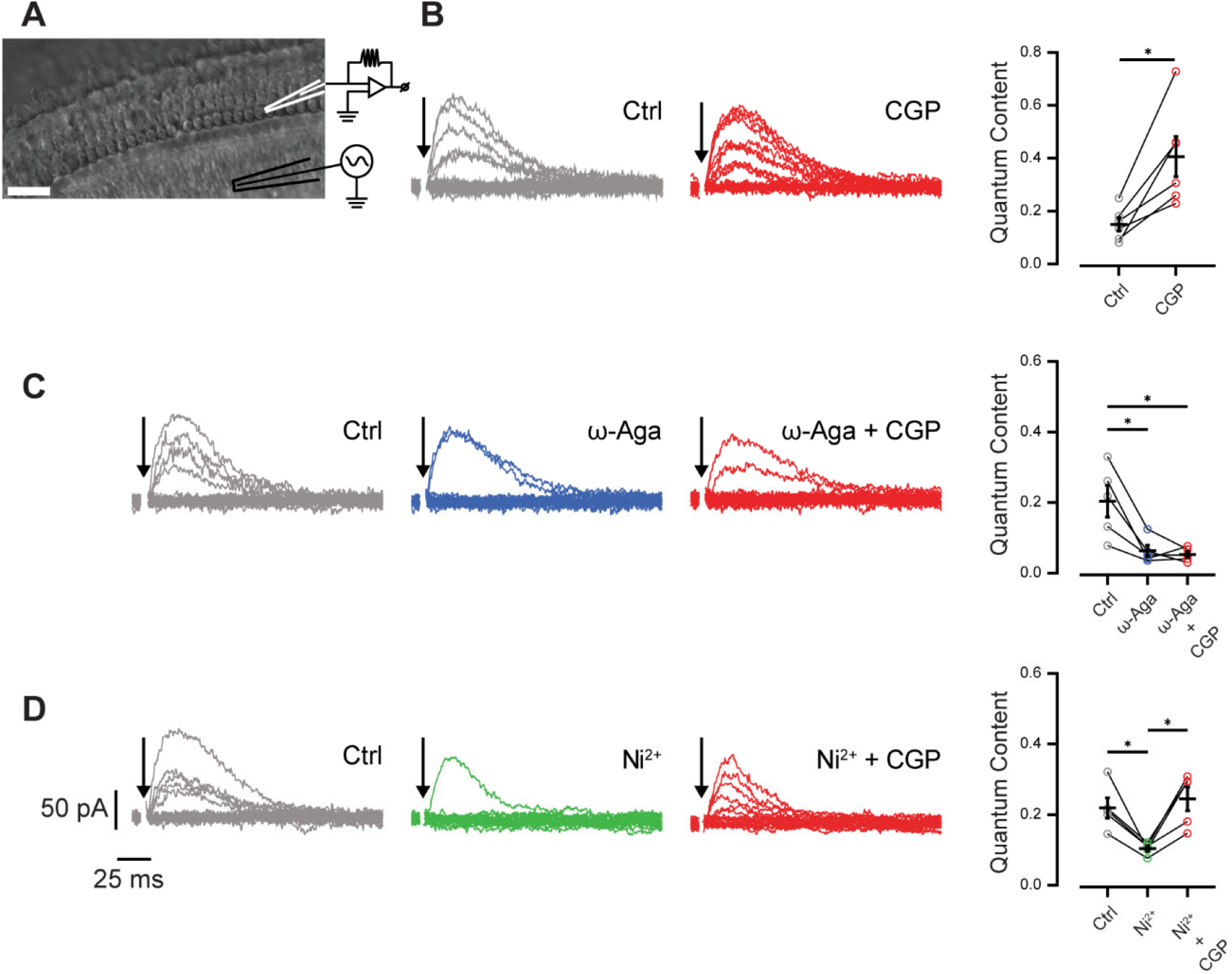
Regulation of GABA release from MOC axons in the OSB by GABA_B_Rs. A. Schematic of the cochlear preparation and the position of the recording and stimulating electrodes. B. Left panels: representative recordings of synaptic responses in P11-13 OHCs in response to electrical stimulation of MOC axons in control conditions and in the presence of the GABA_B_R blocker CGP 36216 (100 µM). Right panel: quantification of quantal content in both conditions. C. Left panels: representative recordings of synaptic responses in control conditions and in the presence of the P/Q-type VGCC blocker ω-agatoxin IVA (200 nM) and in the presence of ω-agatoxin IVA (200 nM) plus CGP 36216 (100 µM). Right panel: quantification of quantal content in the three conditions. D. Left panels: representative recordings of synaptic responses in control conditions, in the presence of the R-type VGCC blocker, Ni^2+^ (100 µM) and in the presence of Ni^2+^ (100 µM) plus CGP 36216 (100 µM). Right panel: quantification of quantal content in the three conditions. Stimulation artifacts were removed for clarity in all traces (arrows).

### MOC axon terminals lack fast GABAergic responses

The above experiments indicate that MOC axons are likely GABAergic and release GABA that acts at presynaptic GABA_B_ autoreceptors. Single-cell transcriptomic experiments indicate GABA_A_R subunits are also expressed in MOC neurons (Frank et al., 2023), while GABA_A_R knockout animals exhibit auditory function and innervation deficits (Maison et al., 2006). These experiments, however, would not be able to distinguish between somatic and axonal GABA_A_R localization, and indeed, MOC somata receive inhibitory GABAergic and glycinergic innervation from the medial nucleus of the trapezoid body (MNTB) (Torres Cadenas et al., 2020, 2022). To directly test whether MOC axon terminals also have fast, ionotropic GABA-evoked responses indicative of GABA_A_R signaling, we performed whole-cell patch-clamp recordings from axons of MOC neurons in cochlear preparations from the apical turn of P13-17 C57Bl/6J WT mice, an age at which we found MOC axons to be accessible for these experiments. A fluorescent tracer (AlexaFluor 488 hydrazide) included in the pipette solution diffused into the cell for *post hoc* identification of the MOC efferent axons based on neuronal morphology (**Figure 3A**). A series of voltage steps was applied to the neuron to assess cell health, and the subsequent voltage-gated conductances confirmed neuron viability. Although voltage-gated conductances were not fully characterized due to the limited number of experiments, recordings suggest the presence of A-type potassium currents, HCN-mediated I_h_ currents, and single action currents (escaping spikes) in response to stimulation (**Figure 3B**). During voltage-clamp recordings (V_hold_ = -80 mV), GABA (1 mM) was applied via a local gravity-driven wide-bore pipette. In four MOC neuron recordings, no currents were evoked by GABA in the MOC axons (**Figure 3C**), indicating that MOC axons and terminals do not have ionotropic GABA_A_R signaling.

**Figure 3.**
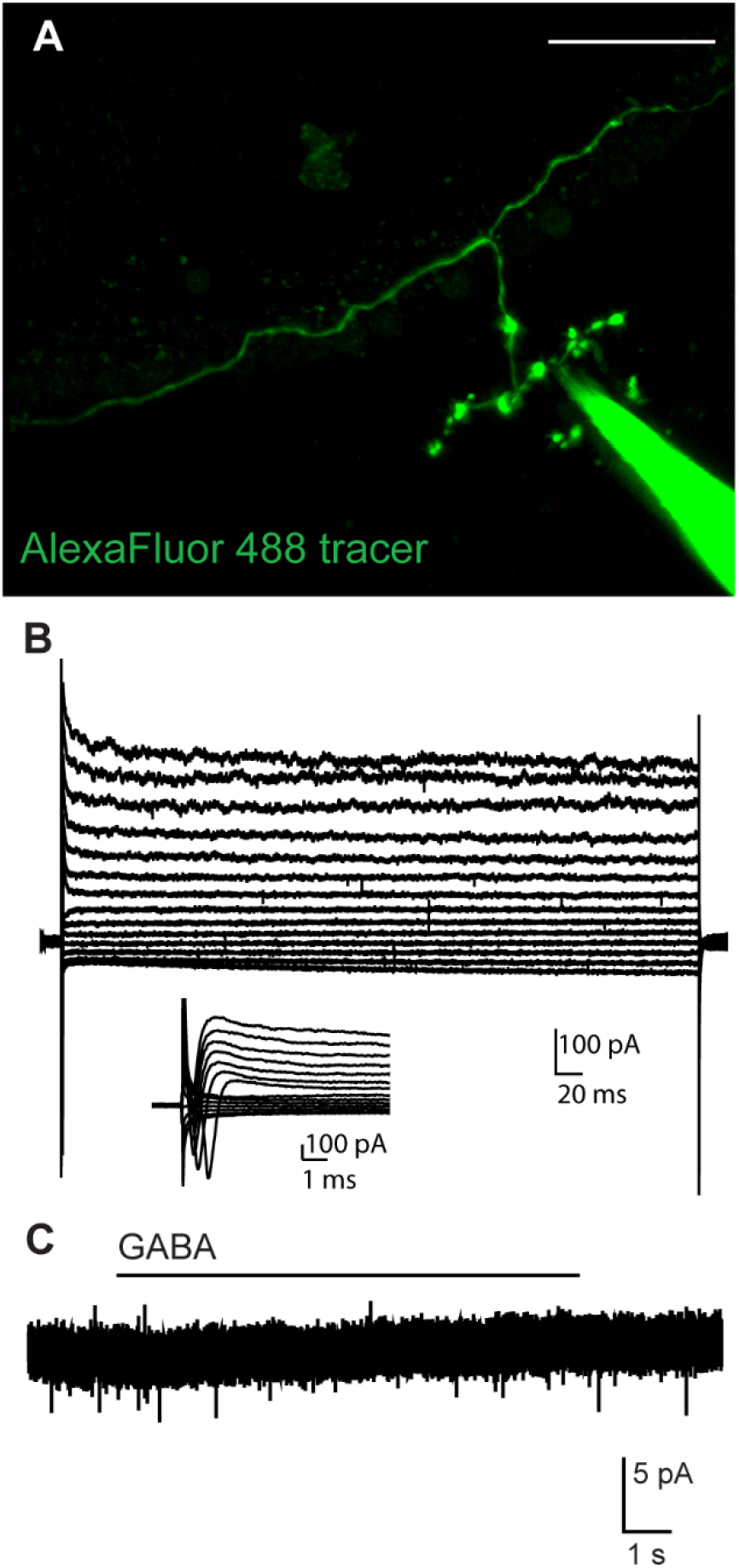
MOC axon terminals lack ionotropic GABA receptors. A. Confocal Z-stack of a single MOC axon terminal from which patch-clamp recordings were performed, filled with AlexaFluor 488 hydrazide diffused from the patch pipette. Scale bar = 50 µm. B. Set of current responses to hyperpolarizing and depolarizing voltage steps in 10 mV increments from -120 to +10 mV in 10 mV increments. Inset shows zoom of action currents from beginning of voltage step traces. C. Whole-cell voltage-clamp trace from the same MOC axon as in (B) during application of 1 mM GABA.

### GABA is released from MOC axons onto type II SGN dendrites

The above work indicates that GABA is synthesized in MOC axon terminals and demonstrates that released GABA acts pre-synaptically through activation of GABA_B_Rs but not GABA_A_Rs. However, it has not yet been shown that GABA released from MOC axon terminals diffuses to a post-synaptic site. To explore this, we combined electrical stimulation of neurotransmitter release from MOC axons with optical detection of the site of extracellular GABA transients in the OSB. To detect GABA, the GABA-sensing fluorescent reporter iGABASnFR (Marvin et al., 2018) was expressed in SGNs. To drive iGABASnFR expression in the SGN, an AAV with a neuron-specific serotype and promotor was injected into the posterior semi-circular canal of P1-2 mice. Type II SGN dendrites were then imaged using confocal microscopy to detect an increase in iGABASnFR fluorescence that indicates binding of GABA. While the imaged fluorescent structures had clear morphological patterns of type II SGN dendrites, it was not possible to differentiate between individual dendrites in imaging experiments; ROIs were selected that likely encompassed multiple fibers.

To verify that iGABASnFR does not respond to ACh, which is the primary neurotransmitter of the MOC efferent system and confirm the specificity of iGABASnFR for detecting GABA, we measured changes in iGABASnFR fluorescence in response to exogenously applied neurotransmitters in P12-15 mouse apical turns. Following baseline imaging of type II SGNs (20-30 sec), we applied either ACh or GABA (1 mM, 20-30 sec) using a large-bore gravity-fed pipette. ACh did not evoke an increase in iGABASnFR fluorescence (ACh application compared to 10 sec prior: ΔF/F = -0.049, n = 39 ROIs across 6 cochlear preparations, negative value due to steady bleaching during the experiment). By contrast, application of GABA evoked a significant increase in iGABASnFR fluorescence (ΔF/F = 0.037, n = 44 ROIs across 6 cochlear preparations, one-way ANOVA, GABA vs. ACh p = 0.0013; **Figure 4A-C**). These experiments confirm that iGABASnFR responds specifically to GABA, but not to ACh.

**Figure 4.**
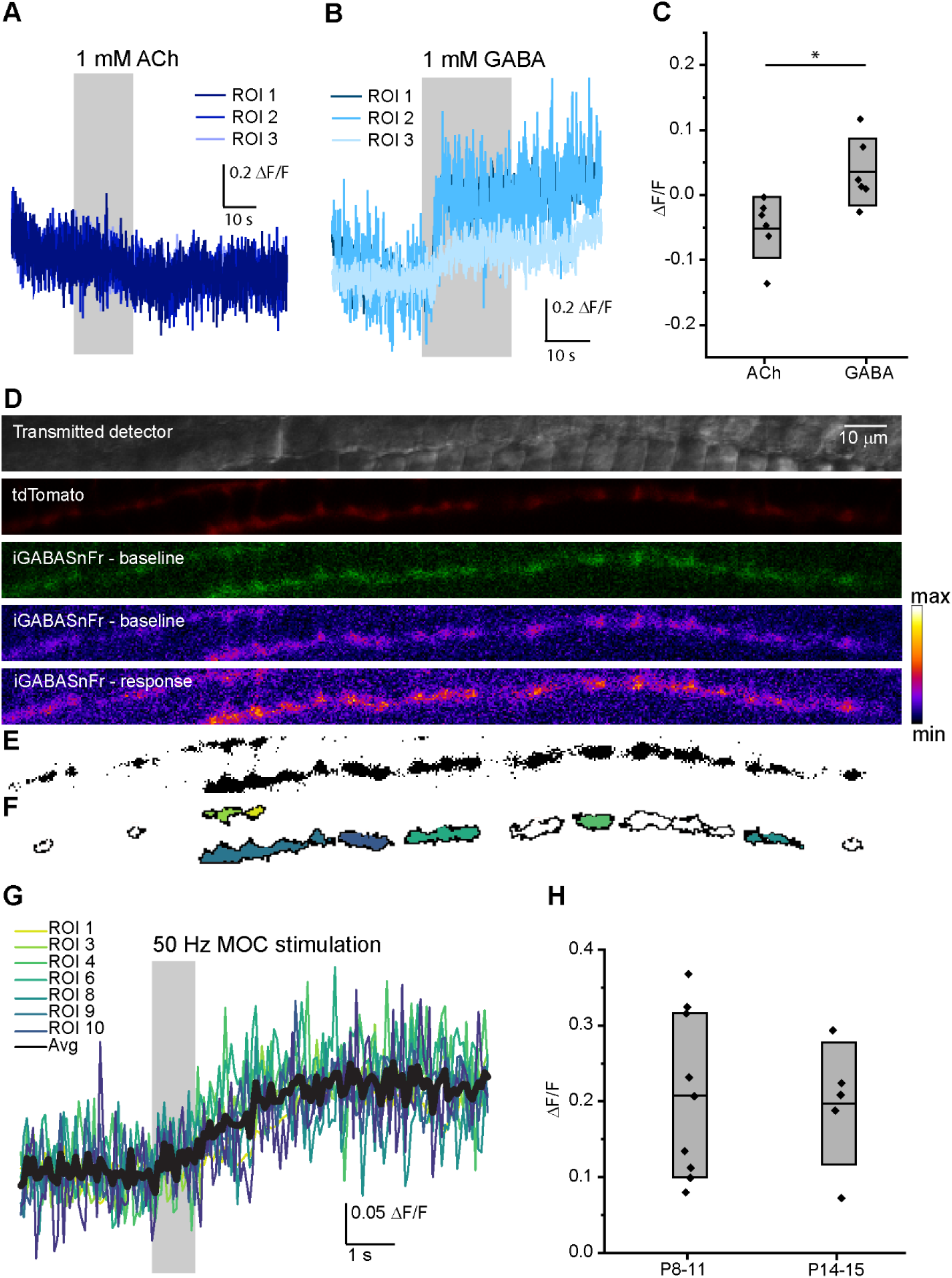
Endogenous release of GABA from MOC axons in the OSB. A. Fluorescence responses from three ROI’s from an example representative experiment with iGABASnFR expressed in type II SGNs, which do not change in response to application of exogenous ACh (1 mM). B. Fluorescence responses from three ROI’s from an example representative experiment with iGABASnFR expressed in type II SGNs, which show an increase in fluorescence in response to application of exogenous GABA (1 mM). C. Quantification of iGABASnFR response to exogenously applied ACh and GABA, each dot represents mean ΔF/F of all ROIs from a single experiment, box indicates mean and SD of population data. D. Example images from a type II SGN transduced with iGABASnFR2.0 from a P8 mouse. The transmitted detector image shows the location of imaging near the base of OHCs. The tdTomato image from a Bhlhb5-Cre; tdT animal shows the location of the type II SGN. The iGABASnFR imaging is visible in baseline (green) images, and then using a heatmap to show fluorescence levels in baseline conditions (“baseline”) and after electrical stimulation of MOC axon terminals (“response”). E. Thresholded image from the maximum intensity projection of the type II SGN shown in ‘D’. F. ROIs used for iGABASnFR analysis. Color coding relates to ROI #’s in (G), white ROI’s indicate regions that did not respond to electrical stimulation with a fluorescence increase. G. ΔF/F plots showing the time-course of iGABASnFR fluorescence before, during, and after electrical stimulation of pre-synaptic MOC axons (50 Hz, 1 sec), at ROIs indicated by colors in (F). H. Summary data for iGABASnFR fluorescence responses in type II SGNs at two post-natal age groups. Each dot represents mean ΔF/F values for a single experiment, box indicates mean and SD.

Next, the activity-dependent release of endogenous GABA from MOC axons in the OSB was tested by stimulating neurotransmitter release from MOC axon terminals while imaging iGABASnFR fluorescence. Wild type (WT) C57Bl/6J mice were used along with Bhlhb5 Cre; tdT and NgnCreERT2; tdT mice that expressed the red fluorescent protein tdTomato in cochlear neurons to allow co-localization of the iGABASnFR with SGNs. The cochlear apical turn from AAV-iGABASnFR injected mice were dissected at P8-11 (before hearing onset, iGABASnFR2.0 FLEX, 6-9 days after injection) or P14-15 (after hearing onset, WT mice with iGABASnFR.F102G, 12-13 days after injection). The type II SGNs were localized under confocal microscopy using either iGABASnFR basal fluorescence or tdTomato fluorescence. Control images were collected for ∼3 sec, followed by electrical stimulation of MOC axons (50 Hz, 1 sec) while imaging continued for an additional ∼6 sec (**Figure 4D-F**). There was a significant increase in iGABASnFR fluorescence in type II SGNs in response to electrical stimulation of MOC axons from both pre-hearing (P8-11, example **Figure 6G**) and post-hearing onset mice (P14-15). Although the change in fluorescence between the two age groups is not directly comparable due to differences in the iGABASnFR variants used and time course of expression, there was no difference in the mean amplitude of responses between the two groups: (P8-11 (iGABASnFR.2 FLEX): mean ΔF/F 0.208 ± 0.109, 1 to 8 positive ROIs per experiment, 9 of 17 experiments had at least one ROI with a positive response; P14-15 (iGABASnFR1.0): mean ΔF/F 0.197 ± 0.080, 1 to 4 ROIs per experiment, 5 of 6 experiments had positive response, one-way ANOVA comparing age groups, p = 0.85; **Figure 4H**). These results support an activity-dependent release of GABA from MOC axon terminals both before and after the onset of hearing. This is consistent with GABA released from MOC axon terminals that acts at pre-synaptic GABA_B_R autoreceptors, as shown in both previously published work (Wedemeyer et al., 2013), and the results above. Notably, the iGABASnFR imaging data shown here demonstrates that GABA also diffuses from MOC terminals to other locations in the OSB, supporting a broader potential role.

### Type II SGN dendrites have functional GABA_A_Rs

To further probe whether GABA diffusion from pre-synaptic MOC axon terminals would act at a post-synaptic cell in the OSB, cochleae were immunolabeled with antibodies against the GABA_A_R β3 subunit. Previous reports indicated GABA_A_R β3 subunit expression in the cochlea, with knockouts causing auditory dysfunction and innervation deficits (Maison et al., 2006). Here, the GABA_A_R β3 localized at punctate structures in the cochlea, specifically in the OSB (**Figure 5A**). GABA_A_R β3 immunoreactivity did not co-localize with GAD-immuno-positive, genetically labeled MOC terminals in ChAT-IRES-Cre; tdT mice, suggesting that MOC terminals do not have this GABA_A_R subunit. This is consistent with above recordings indicating that MOC axon terminals lack GABA_A_Rs. Taking advantage of sparse afferent labeling that occurs in Ngn1CreERT2; tdT mice without tamoxifen treatment (Koundakjian et al., 2007), immunolabeling for GABA_A_R β3 subunits clearly localized to genetically labeled, (tdT-positive) type II SGN terminals as well as the presumptive type II SGN terminals that lacked Ngn1CreERT2 (tdT-negative) (**Figure 5B**). The GABA_A_R β3 antibody-positive dendritic bouton structures were directly opposed to GAD-immuno-positive MOC terminals. This labeling suggests the potential for fast, ionotropic GABAergic responses in type II SGNs.

**Figure 5.**
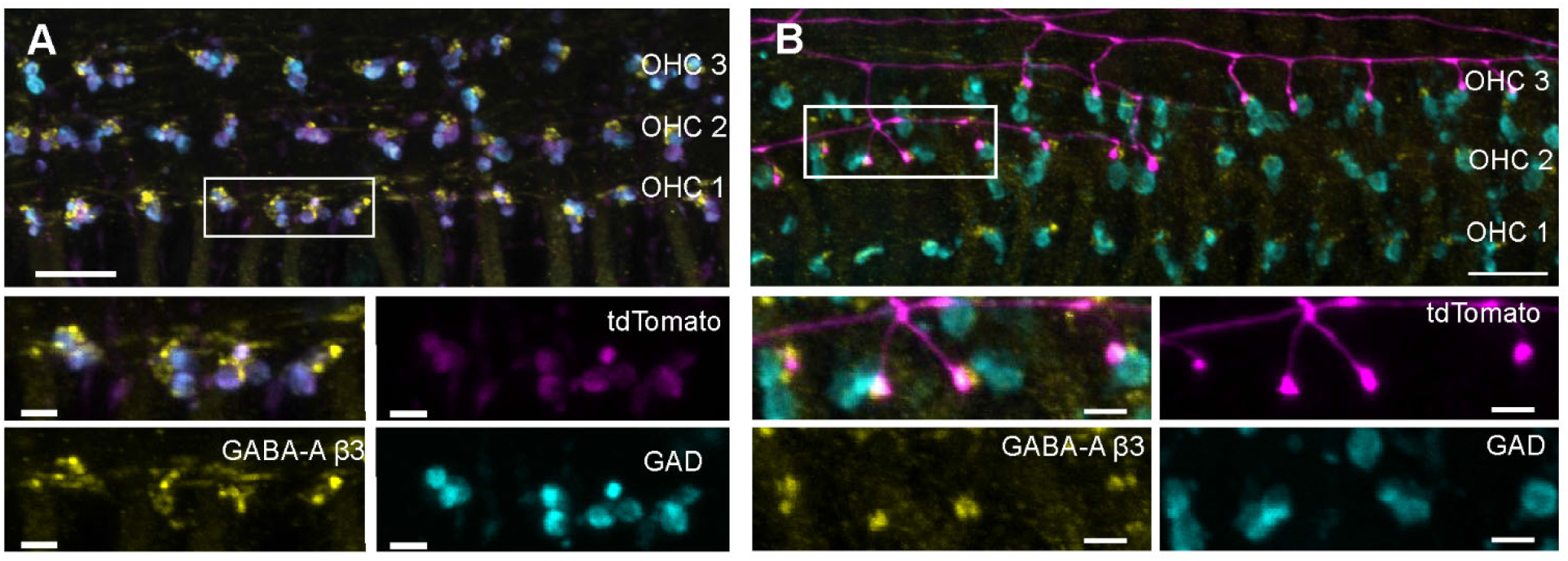
GABA_A_R β3 localized to type II SGNs. A. Immunolabeling of GAD (cyan) in genetically identified MOC axon terminals (magenta) in a P20 ChAT-IRES-Cre; tdT mouse. An anti-GABA_A_R β3 antibody (yellow) labels structures adjacent to MOC terminals and is absent from the MOC terminals themselves. Box in top panel indicates region for zoomed bottom panels. B. Immunolabeling of GAD (cyan), adjacent to genetically labeled type II SGN dendrites in a P20 Ngn1CreERT2; tdT mouse (magenta), which has random, sparse expression of Cre. Immunolabeling of the GABA_A_R β3 (yellow) localizes to boutons in axon branches from tdT-labeled type II SGNs. Additional GABA_A_R β3 labeling in non-red structures is likely also in type II SGNs that were not labeled with the sparse Ngn1CreERT2; tdT expression. Box in top panel indicates region of zoom for bottom panels. Scale bar in top panels = 10 µm, scale bar in zoom panels = 2.5 µm.

Immunolabeling experiments above indicate that type II SGN neurons have GABA_A_Rs near GAD-positive MOC axon terminals, while the above patch-clamp and iGABASnFR experiments demonstrate activity-dependent release of GABA from MOC axon terminals that diffuses to type II SGNs. To further investigate whether type II SGNs directly respond to GABA, whole-cell patch-clamp recordings (Weisz et al., 2009, 2012, 2014) were made from the dendrites of type II SGNs from the apical turns of P5-10 WT C57BL/6J mice. AlexaFluor 488 hydrazide was included in the pipette solution for *post-hoc* identification based on characteristic type II SGN morphology (Weisz et al., 2009, 2012; Liu et al., 2015; Martinez-Monedero et al., 2016) (**Figure 6A, B**). The type II SGN dendrite was held at a membrane voltage of -80 mV to accentuate chloride-mediated currents (ECl = -25 mV). GABA (1 mM) was applied (duration 3-20 sec) via a gravity fed, large-bore application pipette. GABA application evoked currents of -22.6 ± 18.0 pA (n = 31 neurons), which returned to baseline after GABA application ceased (**Figure 6C**). Such GABAergic currents were detected in type II SGN dendrites through P15, the oldest age at which experiments were successfully performed (not shown), although the difficulty of recordings at these older ages precluded extensive pharmacology studies. To test whether GABA-evoked currents were due to activation of ionotropic post-synaptic receptors, the membrane potential was stepped from -80 to +40 mV, in 20 mV increments, followed by GABA application (1 mM, 10-20 sec). GABA-evoked currents reversed polarity at ∼-12 mV (∼-21 after Liquid Junction Potential correction), close to the calculated reversal potential of chloride for the experimental solutions used, supporting neurotransmission through ionotropic GABA_A_Rs (**Figure 6E**). To further test for activation of GABA_A_Rs, GABA was first applied in control conditions, then in the same cells during bath application of the ionotropic GABA_A_R blocker gabazine (SR 95531, 30 µM), followed by a third application of GABA 10-20 min after a wash of gabazine. Gabazine application significantly reduced GABA-evoked currents (control mean: -38.1 ± 23.0 pA, gabazine: -16.1 ± 11.4 pA, wash: -19.3 ± 12.6 pA, one-way ANOVA p = 0.017, n = 8-10 neurons, **Figure 6C, D**), suggesting that GABA-evoked currents are mediated at least in part by ionotropic GABA_A_R in the type II SGN.

**Figure 6.**
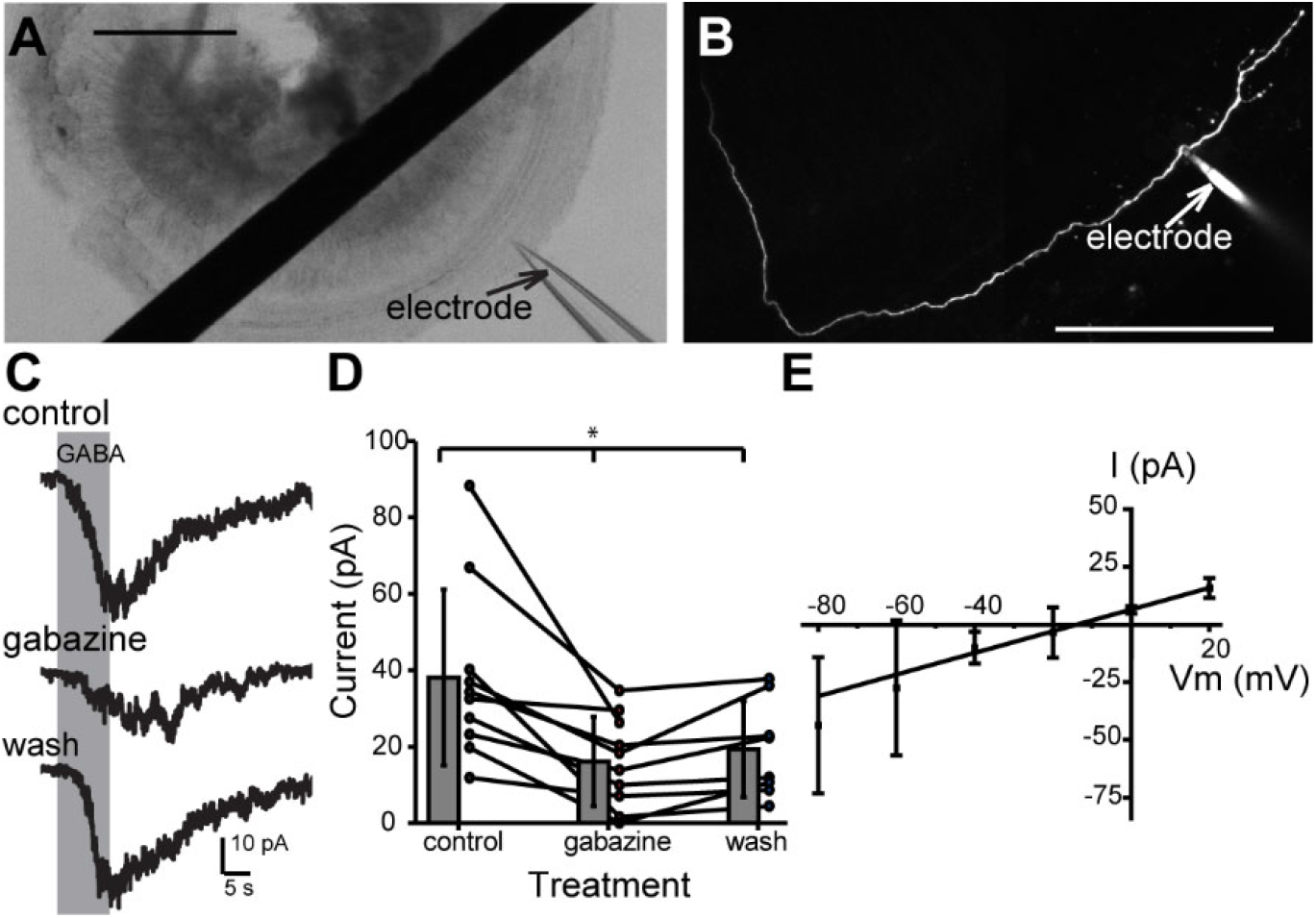
GABA_A_R mediated responses in type II SGNs. A. DIC image of acutely dissected cochlear apical turn during patch-clamp recording from a type II SGN dendrite. Scale bar = 250 µm. B. A single type II SGN dendrite immediately following whole-cell voltage-clamp recordings, filled with AlexaFluor 488 hydrazide via the patch pipette. Scale bar = 100 µm. C. Voltage-clamp recordings from a type II SGN (V_hold_ = -80 mV) during 10 sec focal application of 1 mM GABA in control conditions (top), with 30 µM gabazine in the bath solution (middle), and after wash of gabazine (bottom). D. Summary data showing magnitude of currents evoked by 1 mM GABA application in control, gabazine, and wash conditions. E. Current-voltage relation of 1 mM GABA responses from type II SGN at different holding potentials between -80 mV and +20 mV in 20 mV increments, with linear fit.

Our combined results demonstrate immunolocalization of GABAergic components (GAD, GABA_A_R β3) in genetically identified MOC axons and type II SGN dendrites that are suggestive of a functional synapse. We confirmed that activity-dependent GABA release reduces further activity at pre-synaptic MOC axon terminals by autoreceptor GABA_B_R-mediated inhibition of P/Q type VGCC, adjusting the strength of the efferent synapse. Further, evoked release of GABA from MOC axon terminals also diffuses to post-synaptic sites at the type II SGN. Finally, we recorded GABA_A_R mediated responses in type II SGNs. Together, these results indicate complex roles of GABA in the OSB that include both control of MOC efferent synaptic strength and a GABAergic synapse between MOC efferent neurons and type II SGN afferents (**Figure 7**).

**Figure 7.**
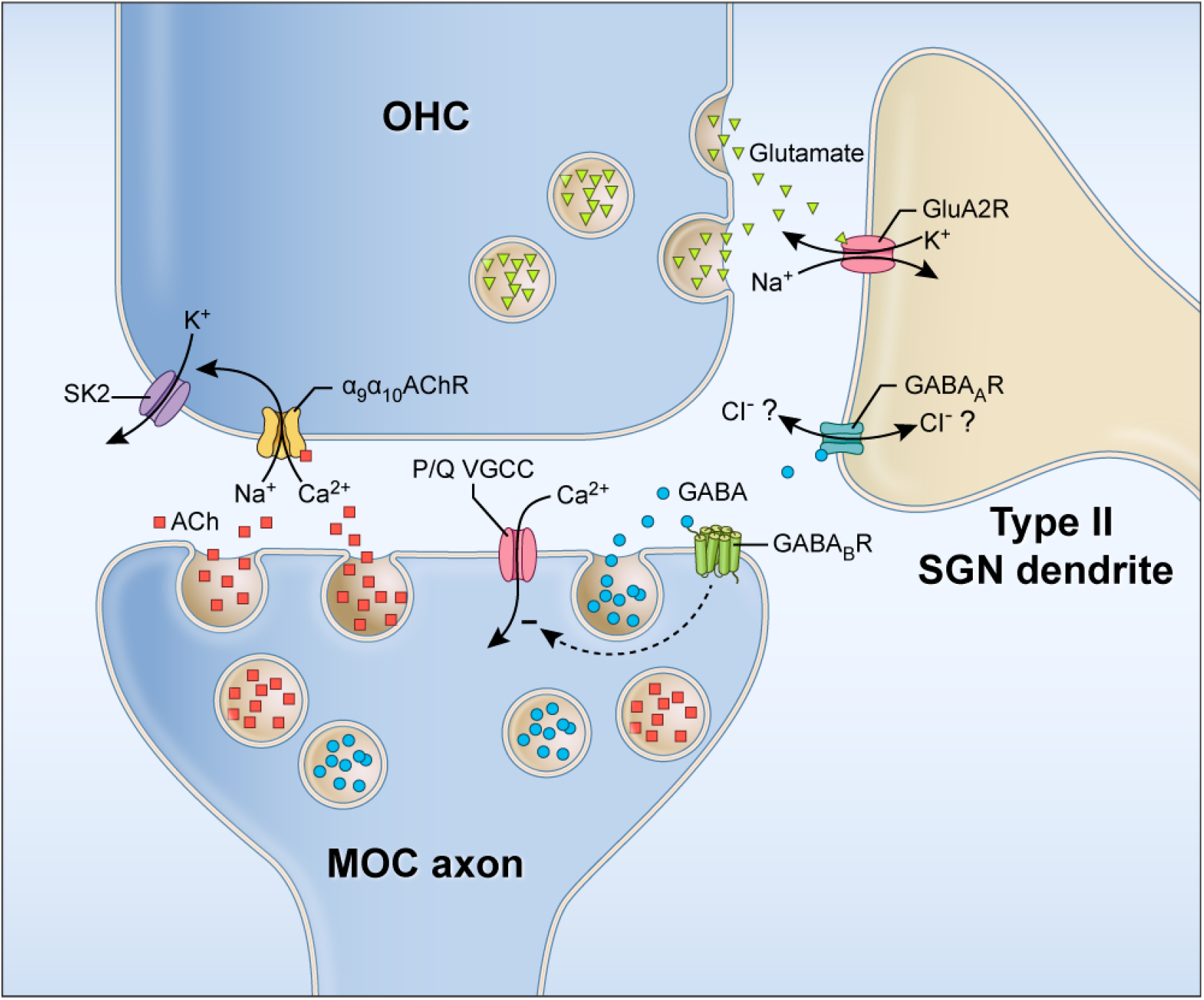
Schematic of neurotransmitter signaling between OHC, type II SGN afferent neurons, and MOC efferent neurons. ACh (red squares) and GABA (blue circles) are released from MOC axons. ACh acts at α_9_/α_10_nAChR on OHCs. Calcium influx through the nAChR activates SK2 potassium channels to hyperpolarize OHCs. Glutamate (green triangles) released from OHCs activates GluA2R on type II SGN dendrites. GABA released from MOC axon terminals acts at pre-synaptic GABA_B_R autoreceptors on the MOC axon terminals to decrease calcium influx through P/Q type VGCC. This then reduces neurotransmitter release from the MOC terminals, disinhibiting OHCs and possibly type II SGNs. GABA also diffuses from MOC terminals to act at GABA_A_Rs on type II SGN dendrites, although the effect on the type II SGN dendrite will depend on the chloride concentration gradient between the type II SGN dendrites and the extracellular space, as suggested by the double-headed arrow indicating that the direction of Cl^-^ flow through the GABA_A_R is unknown. Schematic by Alan Hoofring, NIH Medical Arts Branch.

## Discussion

The mammalian cochlea uses fast, high-fidelity synapses and 1:1 innervation between IHCs, the primary auditory sensory receptors, and type I SGN afferents to respond to sound stimuli with temporal precision and accurate frequency encoding. The more numerous, electromotile OHCs are also critical for normal hearing due to their ‘cochlear amplifier’ functionality that lowers the threshold for sound detection and improves frequency tuning of the cochlea. The MOC efferent system inhibits OHC activity, thus decreasing the OHC-mediated enhancement of basilar membrane vibrations. This MOC activity is implicated in reduced responses to sustained background noises that enable detection of salient sounds, protection of the cochlea against acoustic overstimulation, and auditory attention (reviewed in Fuchs and Lauer, 2019). Although the functions of the MOC and OHC systems are well documented, the roles of other cell types and potential circuits in this region remain elusive. For example, OHC afferent innervation via the small population of type II SGN afferents has been clearly demonstrated, yet the role of this system in hearing is unclear (reviewed in Fuchs and Glowatzki, 2015). Furthermore, in addition to the canonical MOC-OHC and OHC-type II SGN circuitry in the OSB, ultrastructural studies suggest numerous additional synaptic contacts including among OHCs, MOC efferents, type II SGN afferents, and supporting cells (Eybalin et al., 1988; Burgess et al., 1997; Fechner et al., 1998, 2001; Thiers et al., 2002a, 2002b, 2008; Hua et al., 2021). Whether these putative synapses are active in hearing as well as which neurotransmitters are involved, has not yet been determined. As a first step to understanding how intricate circuits in the OSB contribute to hearing, we used patch-clamp electrophysiology recordings, immunolabeling, and optical neurotransmitter detection to demonstrate activity at one of these putative synapses, between MOC efferent and type II SGN afferent neurons in the OSB, that we propose is GABAergic.

The potential role of a GABAergic MOC to type II SGN synapse could include both central and peripheral mechanisms. Although OHC release of glutamate onto type II SGNs is sparse and weak (Weisz et al., 2009, 2012), glutamatergic OHC-type II SGN signaling centrally to the cochlear nucleus granule cell domains has been demonstrated in response to loud, but non-damaging, broadband sound (Weisz et al., 2021). MOC neurons also send axon branches to the granule cell domains (Brown et al., 1988; Beebe et al., 2023), suggesting multiple sites of interaction between MOC efferent and type II SGN afferent systems that may report MOC activity back to the brain for fine-tuning of efferent activity. While the contributions of a GABAergic MOC-type II SGN synapse to central signaling are yet to be determined, these synapses exemplify another direct interaction between afferent and efferent cochlear neurons.

It is also possible that the MOC-type II SGN synapses have a role in peripheral cochlear function. Given the robust hyperpolarizing response of OHCs to MOC-mediated cholinergic transmission, however, it is perplexing why MOC neurons would then provide additional direct GABAergic inhibition of the type II SGN afferents. This is unexpected for a signaling pathway (the OHC to type II SGN pathway) that is very weak. That stated, we do not yet know whether activation of GABA_A_Rs on type II SGN under physiological conditions is indeed inhibitory, due to the possibility of a relatively depolarized Cl^-^ reversal potential that is common in the developing nervous system and may persist in dendritic or axonal processes of mature neurons (Turecek and Trussell, 2001; Alle and Geiger, 2007; Stell et al., 2007; Sernagor et al., 2010; Pugh and Jahr, 2011; Weisz et al., 2016). Immunohistochemistry studies shown here with genetically labeled efferents demonstrate the GABAergic nature of MOC terminals prior to hearing onset, before MOC terminals have achieved their mature innervation patterns and when neurons are more likely to have a depolarizing chloride reversal potential. Due to limitations in dendritic recording techniques, patch-clamp recordings of GABA_A_R -mediated activity in type II SGNs were primarily performed prior to P11 and included an artificial chloride concentration; the actual concentration has not been determined. Thus, GABAergic signaling may be excitatory in developing cochlear neurons to support developmental roles such as synapse formation, while the polarity of GABAergic responses in mature SGNs is at this time unknown. Regardless, the aforementioned recording experiments indicate that type II SGNs do respond to GABA prior to hearing onset, while results from iGABASnFR imaging of activity-dependent GABA release and immuno-labeling of GABAergic structures, suggest that GABAergic MOC to type II SGN signaling also occurs in the mature cochlea.

The pre- and post-synaptic functions of GABA in the OSB contribute additional complexity. GABA acts as a negative regulator of MOC axon terminal function, reducing ACh release by acting at pre-synaptic GABA_B_R. Here we show that this activity occurs via the same mechanism as at MOC-IHC synapses, by GABA_B_R-mediated decreases in pre-synaptic P/Q VGCC activity. Therefore, while ACh activity at OHCs leads to SK channel-mediated inhibition of OHCs that likely reduces OHC-based activation of type II SGNs, GABA may be simultaneously reducing release of ACh, and likely GABA, from MOC terminals, thus disinhibiting the OHC-type II SGN afferent circuit. It must be borne in mind, however, that we do not yet know whether GABA and ACh are co-released by the same MOC fibers and whether the MOC synaptic terminals reaching type II SGNs are the same and/or have the same complement of presynaptic proteins involved in release and activity-dependence as those contacting the OHCs. Further adding to this complexity, evidence suggests that pre-synaptic GABA_B_R expression is likely downregulated at MOC-OHC synapses following hearing onset because it is no longer observed in the adult (Maison et al., 2009) while in post-synaptic compartments both GABA_A_R (present work; Maison et al., 2006) and GABA_B_R expression (Maison et al., 2009) is maintained in the adult. The functional interplay of pre- vs. post-synaptic and receptor subtype-specific GABA signaling over the course of development in both potentially decreasing and/or increasing type II SGN activity indicates that additional study into the particular patterns and conditions of GABA transmission is necessary.

In summary, our results indicate that there is a functional, GABAergic synapse between MOC efferent and type II SGN afferent neurons, which operates via post-synaptic GABA_A_R. While the role of this MOC-type II SGN synapse is unclear, this is the first confirmation that one of the abundant complex circuits in the OSB previously identified by electron microscopy is indeed functional. While future work will delineate the role of this synapse in cochlear function, the present results indicate that function of the OSB, which is critical for fine-tuning cochlear activity and gain control, is likely more complex than previously demonstrated.

## Acknowledgements

Thank you to the GENIE Project Team at Janelia Research Campus, HHMI for developing the iGABASnFR 1 and 2 variants, and for providing iGABASnFR2 plasmids. Thank you to Dr. Matthew Kelley and Dr. Katie Kindt for critical readings of an earlier draft of the manuscript. The synapse schematic is credited to Alan Hoofring, NIH Medical Arts Branch. This research was in part supported by the Intramural Research Program of the NIH, NIDCD, Z01 DC000091 (CJCW), and in part supported by Agencia Nacional de Promoción Científica y Técnica Argentina (PICT 2018-02531 CW and PICT 2018-01189 EK).

